# The relationship between spectral and plant diversity: disentangling the influence of metrics and habitat types

**DOI:** 10.1101/2022.09.05.506583

**Authors:** Perrone Michela, Di Febbraro Mirko, Conti Luisa, Divíšek Jan, Chytrý Milan, Keil Petr, Carranza Maria Laura, Rocchini Duccio, Torresani Michele, Moudrý Vítězslav, Šímová Petra, Prajzlerová Dominika, Müllerová Jana, Wild Jan, Malavasi Marco

## Abstract

Biodiversity monitoring is crucial for ecosystem conservation, yet field data collection is limited by costs, time, and extent. Remote sensing represents a convenient approach providing frequent, near-real-time information over wide areas. According to the Spectral Variation Hypothesis (SVH), spectral diversity (SD) is an effective proxy of environmental heterogeneity, which ultimately relates to plant diversity. So far, studies testing the relationship between SD and biodiversity have reported contradictory findings, calling for a thorough investigation of the key factors (e.g., metrics applied, ecosystem type) and the conditions under which such a relationship holds true. This study investigates the applicability of the SVH for plant diversity monitoring at the landscape scale by comparing the performance of three different types of SD metrics. Species richness and functional diversity were calculated for more than 2000 cells forming a grid covering the Czech Republic. Within each cell, we quantified SD using a Landsat-8 “greenest pixel” composite by applying: i) the standard deviation of NDVI, ii) Rao’s Q entropy index, and iii) richness of “spectral communities”. Habitat type (i.e., land cover) was included in the models describing the relationship between SD and ground biodiversity. Both species richness and functional diversity show positive and significant relationships with each SD metric tested. However, SD alone accounts for a small fraction of the deviance explained by the models. Furthermore, the strength of the relationship depends significantly on habitat type and is highest in natural transitional areas. Our results underline that, despite the stability in the significance of the link between SD and plant diversity at this scale, the applicability of SD for biodiversity monitoring is context-dependent and the factors mediating such a relationship must be carefully considered to avoid drawing misleading conclusions.

**Highlights:**

- Plant species richness and functional diversity show significant and positive relationships with spectral diversity
- Spectral diversity alone explains a small fraction of the total variability in ground biodiversity
- Slight differences among the performances of the spectral diversity metrics tested
- The relationship between spectral and plant diversity is context-dependent

## 1. Introduction

Biodiversity supports multiple ecosystem functions, which ultimately provide the ecosystem services essential to sustain human societies (Cardinale et al., 2012; de Groot et al., 2002). However, human exploitation of Earth’s natural resources has led to alterations in plant species’ distribution, composition, and abundance, to the extent that over one-fifth of all vascular plant species are threatened (Willis, 2017). Nowadays there is an urgent need to improve ways to effectively monitor biodiversity across broad spatial scales and to assess how plant communities respond to global change. Nonetheless, the ability to measure and monitor biodiversity continues to lag far behind (Skidmore et al., 2021).

Remote sensing (hereafter RS) offers remarkable opportunities for biodiversity monitoring, from local to global scales (Rocchini et al., 2010; Wang and Gamon, 2019). Among the approaches studied, spectral diversity (SD), originally developed in the framework of the spectral variation hypothesis (SVH, Palmer et al., 2000, 2002), has been gaining momentum as a basis for relating the remotely sensed spectral signal to ground biodiversity at different spatial scales and resolutions (Fassnacht et al., 2022; Rocchini et al., 2021, 2010; Tagliabue et al., 2020; Torresani et al., 2019). At broader spatial resolution, the underlying assumption is that the spatial variation in reflectance values in a given area (i.e., SD) is likely correlated with the spatial variation in the environment in that area and therefore related to the number of species present. At very fine spatial resolution, the relationship is assumed to be direct, with SD directly related to plant diversity through the diversity of species-specific optical traits associated with plant functional and structural properties (Ustin and Gamon, 2010).

Despite the potential of RS in biodiversity monitoring systems, issues remain regarding the hypothesised relationship between spectral diversity and plant ground diversity (Fassnacht et al., 2022). Although several empirical studies have validated the use of spectral diversity to estimate plant species diversity (Levin et al., 2007; Rocchini, 2007; Rocchini et al., 2014), others have criticised it for being unstable and not reliable in every context (Conti et al., 2021; Schmidtlein and Fassnacht, 2017). Such inconsistent findings may be due to a lack of systematic consideration of the key factors that influence the relationship between plant biodiversity and SD, which complicates the interpretation of spectral variation in many contexts (Fassnacht et al., 2022). Recently, Fassnacht et al. (2022) discussed four of the most important factors that may affect such a relationship: 1) the scale considered, both in terms of spatial extent (i.e., size of the study area) and spatial grain (i.e., pixel size); 2) reflectance changes over time (e.g., seasonality); 3) effects of the metric chosen for quantifying SD; and 4) the identity and number of habitat or vegetation types considered (Rocchini et al., 2018, 2010; Rossi et al., 2021; Schmidtlein and Fassnacht, 2017).

Regarding the latter factor, habitat structure may strongly influence the relationship between SD and biodiversity, so all other factors must be adjusted accordingly. For instance, small patches of calcareous grassland embedded in intensively managed agricultural landscapes are very rich in species. However, when observed with medium resolution sensors, they have low SD, while surrounding agricultural land has high SD but very low species richness (Fassnacht et al., 2022). Thus, not considering the variety and types of habitat can be particularly problematic when multiple habitat types are examined simultaneously, as is the case at large to medium observation scales (Fassnacht et al., 2022; Schmidtlein and Fassnacht, 2017).

Similarly, the SD metrics are thought to be linked to plant diversity through different pathways, which may lead to different results. To date, several SD metrics have been proposed as proxies for biodiversity, with no consensus on which metric best fulfils this role. Wang and Gamon (2019) have described three main classes of SD metrics:

i. metrics based on variation in traditional vegetation reflectance indices, based on the well-known correlation between productivity and plant diversity (Tilman et al., 2001; Zhang et al., 2012);
ii. metrics based on spectral information content, which condense the full-spectral information through statistical metrics of variability or spectral entropy;
iii. metrics based on “spectral species” (or “spectral types”) that share similar spectral signatures and are derived from partitioning the spectral space.

To date, the existing empirical studies investigating the relationship between SD and plant diversity have relied mainly on limited datasets (Fassnacht et al., 2022). In addition, most efforts to link plant diversity at the landscape scale to SD have focused on taxonomic diversity (e.g., species richness or evenness), yet species influence ecosystem functions through their functional composition, diversity, and abundance (de Bello et al., 2010; Tilman et al., 1997). Functional diversity represents the variability in plant functional traits in a given area and is considered an essential component of biodiversity that determines ecosystem processes and stability. The spatiotemporal mapping of functional diversity at large scales could thus support assessing how environmental changes affect ecosystem functioning. Nonetheless, the potential of SD to infer functional diversity at the landscape scale remains to be demonstrated (Cavender-Bares et al., 2022; Frye et al., 2021; Schweiger et al., 2018).

Here we investigate whether plant diversity (species richness and functional diversity) at the landscape scale is related to SD. To assess the context-dependence of such a relationship, we will test the performance of the three types of SD metrics, taking into account the effects of the habitat types. Specifically, we rely on an extensive, spatially continuous dataset of field-collected data covering the Czech Republic and a multi-temporal Landsat-8 OLI composite.

## 2. Materials and Methods

### 2.1 Plant diversity data

Data on plant diversity were obtained from the Pladias Database of the Czech Flora and Vegetation (Chytrý et al., 2021). This nationwide database contains approximately 13 million records on species occurrence (many of them critically reviewed and validated) and traits of almost 5 thousand taxa (species, subspecies, varieties, and hybrids) of the Czech vascular flora (Chytrý et al., 2021; Wild et al., 2019). We used the latest update of taxonomic concepts and nomenclature (Kaplan et al., 2019). More than 60% of the records used for our analysis were collected in the last 20 years (Wild et al., 2019) and include all native and spontaneously established alien vascular plant taxa, as well as some commonly cultivated crops and alien woody plants. The study area covers an altitudinal range from 115 to 1603 m a.s.l. with annual precipitation varying from about 400 mm to 1,450 mm (Brázdil et al., 2021) and a wide range of environmental conditions, from dry and warm forest-steppe areas to mountainous areas with subalpine vegetation (Chytrý et al., 2017). We counted species richness and calculated functional diversity in grid cells of 5’ × 3’ (approx. 6.0 × 5.5 km), forming a grid of 2551 units covering the entire Czech Republic (Figure 1). All marginal grid cells that were not entirely within the country were excluded from our analysis to avoid potential bias from undersampled units.

**Figure 1.**
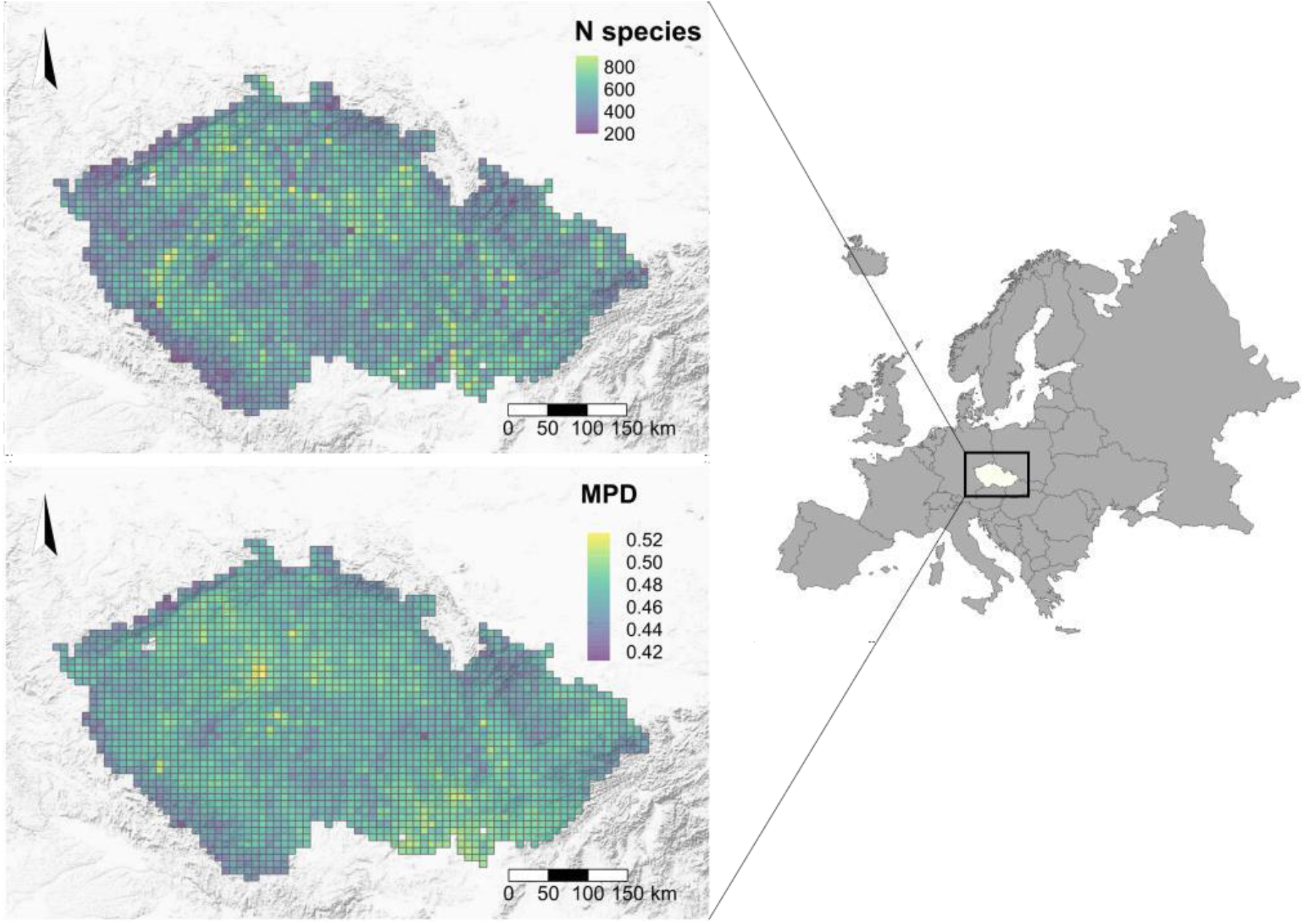
Plant species richness (number of species, top) and functional diversity expressed as mean pairwise distance in plant traits (MPD, bottom) within each grid cell (n = 2363) covering the Czech Republic.

To calculate functional diversity, we selected a list of optically and ecologically relevant functional traits. The majority of traits were available in Pladias (see Chytrý et al. 2021 for their description): Mean Height, Growth Form, Life Form, Leaf Shape, Flower Colour, and three traits were taken to Pladias from the BiolFlor database (Klotz et al. 2002): Leaf Life Span, Leaf Anatomy and Reproduction Type. We also added three traits from the LEDA database (Kleyer et al., 2008) that are not included in the Pladias Database: Specific Leaf Area (SLA), Leaf Dry Matter Content (LDMC), and Seed Mass (Appendix A). We compiled trait values for a minimum of 1463 species for LDMC, and for a maximum of 3087 species for Leaf Shape. To account for potential bias due to missing values, only those grid cells that had at least 70% coverage for each of the traits considered were included in the analysis, resulting in a total of 2363 grid cells. For each grid cell, we calculated functional diversity as the mean pairwise distance (MPD) between species in the functional trait space defined by the above-mentioned traits (Figure 1). Trait distances were measured using Gower distance, where each numerical trait is first standardised by dividing each value by the range of the corresponding trait, after subtracting the minimum value; consequently, the rescaled trait variable has the range [0,1] (Gower, 1971).

### 2.2 Spectral data

We used the USGS Landsat 8 atmospherically corrected Surface Reflectance catalogue of data acquired with the Operational Land Imager (OLI) sensor and available in Google Earth Engine (GEE) (Gorelick et al., 2017). Specifically, we used Earth’s reflected radiance in six bands in the 520–2300 nm range of the electromagnetic spectrum, with a spatial resolution of 30 m and a revisit time of 16 days. To select reflectance data for the same phenological period in which the ground biodiversity measurements were made (i.e., the growing season), we processed OLI images acquired over the area of interest during the growing seasons (i.e., between May and August each year) from 2013 to 2017. The annual range was chosen to overlap with the period of field records. Despite the shorter period covered by OLI than Landsat-7 Enhanced Thematic Mapper Plus (ETM+), we chose the OLI dataset due to the Scan Line Corrector (SLC) ETM+ failure that occurred in 2003, which has affected data acquisition since then.

To minimise weather-related noise, cloud-covered pixels were filtered out based on the Pixel quality attributes (pixel_qa) band (Foga et al., 2017). Then, the Normalized Difference Vegetation Index (NDVI) was calculated and the NDVI layer was added to each image. Finally, we generated a “greenest-pixel” composite image by selecting the pixels with the highest NDVI value of the overlapping images for the studied time period. This was done to best capture the vegetation cover and smooth out inter-annual variability. By selecting the greenest pixel over the entire period, we aimed to get the optimal vegetation cover of the area and avoid the effects of extreme events, such as floods or fires. Another reason for using the greenest-pixel composite is that the relationship between reflectance variability in space and vegetation tends to be higher when it is near the vegetation optimum, i.e., when the cover of photosynthetically active vegetation is greatest (Feilhauer and Schmidtlein, 2011; Thornley et al., 2022).

### 2.3 Spectral diversity metrics

We quantified the SD values within each grid cell using three different methods, each representing one of the main categories proposed by Wang and Gamon (2019): i) the standard deviation of NDVI (sdNDVI), ii) Rao’s Q quadratic entropy index (Rocchini et al., 2017), and iii) richness of “spectral communities” (SpecCom) (Rocchini et al., 2021).

#### 2.3.1 sdNDVI

The Normalized Difference Vegetation Index (NDVI) is a traditionally used vegetation reflectance index positively correlated with primary production (Gillespie et al., 2008). sdNDVI has often been used as a continuous measure of the dispersion (variation) of NDVI values in a given area because it explains a reasonable portion of the variability in the in-situ diversity data (Gillespie, 2005; Gillespie et al., 2009; Gould, 2000; Hall et al., 2010; Levin et al., 2007).

#### 2.3.2 Rao’s Q

Rao’s Q index is a continuous metric that quantifies the difference in reflectance values between two pixels drawn randomly with replacement from a defined set of neighbouring pixels, taking into account their abundance and the relative distance between them:

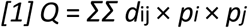

where *d*_ij_ is the spectral distance between pixels *i* and *j*, and p_*i*_ and p_*j*_ are the relative proportions of pixels *i* and *j*, respectively (Rocchini et al., 2017). The advantage of applying Rao’s Q index to spectral data, as opposed to other diversity indices (e.g., Shannon’s H’), is its ability to explicitly account for the numerical size of pixels, rather than just the relative abundance (evenness) of reflectance values. To calculate Rao’s Q index within each grid cell, we used the function `RaoQArea` written in the R language (R Core Team, 2021) and stored in the GitHub repository https://github.com/micheletorresani/RaoQarea. `RaoQArea` calculates Rao’s Q heterogeneity index for a limited area based on the Euclidean distance between pixels of a single-band raster. In our case, the `RaoQArea` function was applied to the single-band raster derived by selecting the first component from a principal component analysis (PCA) applied to a large random subset of pixels of the original multi-band image. PCA was performed to simplify spectral information and remove band collinearity.

#### 2.3.3 SpecCom

To differentiate spectral communities as a function of the optical traits underlying the reflectance of each pixel, we used the R package `biodivMapR` (Féret and de Boissieu, 2022, 2020), which allows spectral diversity mapping based on the partitioning of the spectral space of RS images into subunits called as “spectral species” (following Féret and Asner, 2014). Due to the resolution of the spectral data used, a direct link between the identified spectral subunits and in-situ plant species is not feasible. Therefore, we refer to such partitions in the spectral space as “spectral communities” (*sensu* Rocchini et al., 2021), assuming that they are linked to a higher level of ecological organisation. After a series of pre-processing steps that are part of the package workflow, e.g., spectral normalisation (i.e., continuum removal) and dimensionality reduction (by PCA), the mapping of spectral communities was based on k-means clustering. Clustering was performed on relevant PCA axes selected by visual inspection (Féret and Asner, 2014). The number of k clusters was determined *a priori* and set to 200 after a trial-and-error procedure (Féret and de Boissieu, 2020; Rocchini et al., 2021). Based on the resulting map of “spectral communities”, we calculated the categorical metric “spectral communities” richness (SpecCom) within each grid cell.

### 2.4 Accounting for habitat type

We hypothesised that habitat type may play a role in the relationship between SD and plant diversity (Schmidtlein and Fassnacht, 2017). In the present study, habitat type was defined by CORINE land cover (CLC) because of the broad coverage and validation of these data. We calculated the most represented CLC type (reference year 2012) within each 5’ × 3’ grid cell (i.e., landscape). Although we know that classifying the grid cell with the predominant CLC within the area is an approximation, no other options were possible due to the grain mismatch between the available ground and CLC data. Level I CLC nomenclature was used for artificial land cover types. For forests and semi-natural areas, we assigned land cover codes according to the Level III CLC nomenclature. For agricultural land, we used the Level II nomenclature for arable land and level III for pastures. As a result, the following categories were assigned to the grid cells: CLC 1, CLC 21, CLC 231, CLC 311, CLC 312, CLC 313, CLC 321, and CLC 324 (see Appendix B for a description of the CLC types used). Since CLC 321 (grasslands) was underrepresented (n grid cells= 3), we excluded the corresponding units from the analyses. No landscape features were masked from our spectral data, as this would be in contrast with the original formulation of SVH.

### 2.5 Statistical Analyses

All statistical analyses were performed in R 4.1.2 (R Development Core Team 2021). We modelled the relationship between in-situ diversity and SD through generalised additive models (GAMs) using the mgcv package v. 1.8.39 (Wood 2017), with species richness (SR) or functional diversity (FD) of plants as *response variables* for a total of six models. For species richness, we used a negative binomial (NB) error distribution to account for possible over-dispersed species richness count data. For functional diversity, we used the Gaussian error distribution. Each model had the following three sets of *predictors terms*:

1. An interaction between SD and the dominant CLC type. This represents the effect of CLC type on the relationship between SD and taxonomic or functional diversity. In other words, this models the possibility of different SD-SR or SD-FD relationships in different CLC types.
2. Sampling effort, represented by the logarithm of the number of records within each grid cell. This corrects for the possible effects of uneven sampling effort, i.e., when grid cells may have higher species diversity simply because they have been sampled more.
3. Smooth two-dimensional splines on a sphere (Wood 2017) to account for spatial autocorrelation (i.e. spatial pseudo-replication) in the response variable (Dormann et al., 2007).

To account for possible non-linear patterns, the response of SR and FD to all continuous predictors was modelled as a second-order polynomial. An example of a full model formula as used by the mgcv R package is : `gam(biodiv ∼ poly(SD, 2, raw = F):CLC + s(longitude, latitude, bs = “sos”)+ log(samp.eff`.).

To estimate the relative importance of effects of the three sets of predictors above, we used partitioning of deviance (Aragón et al. 2010; Carrete et al. 2007), an approach related to variance partitioning (Borcard et al. 1992). Specifically, the deviance from a null model with no predictors was partitioned to (i) a fraction explained by spectral diversity and its interaction with CLC, (ii) an effect of sampling effort, (iii) a fraction explained by spatial autocorrelation. We estimated both the independent effects of these, as well as their overlapping fractions, where the overlap is caused by collinearity between the predictors.

The data and code used in the analyses are available at https://github.com/MichelaPerrone/SVH_CZ.git under CC-BY license.

## 3. Results

We found statistically significant relationships between plant diversity (species richness and functional diversity) and SD. The models with species richness and functional diversity as responses were similar, all showing positive relationships with SD. The explanatory power of the models, expressed as explained deviance, ranged from 71.9% to 75.1% for models explaining species richness and from 58.0% to 64.8% for models explaining functional diversity (Figure 2).

**Figure 2.**
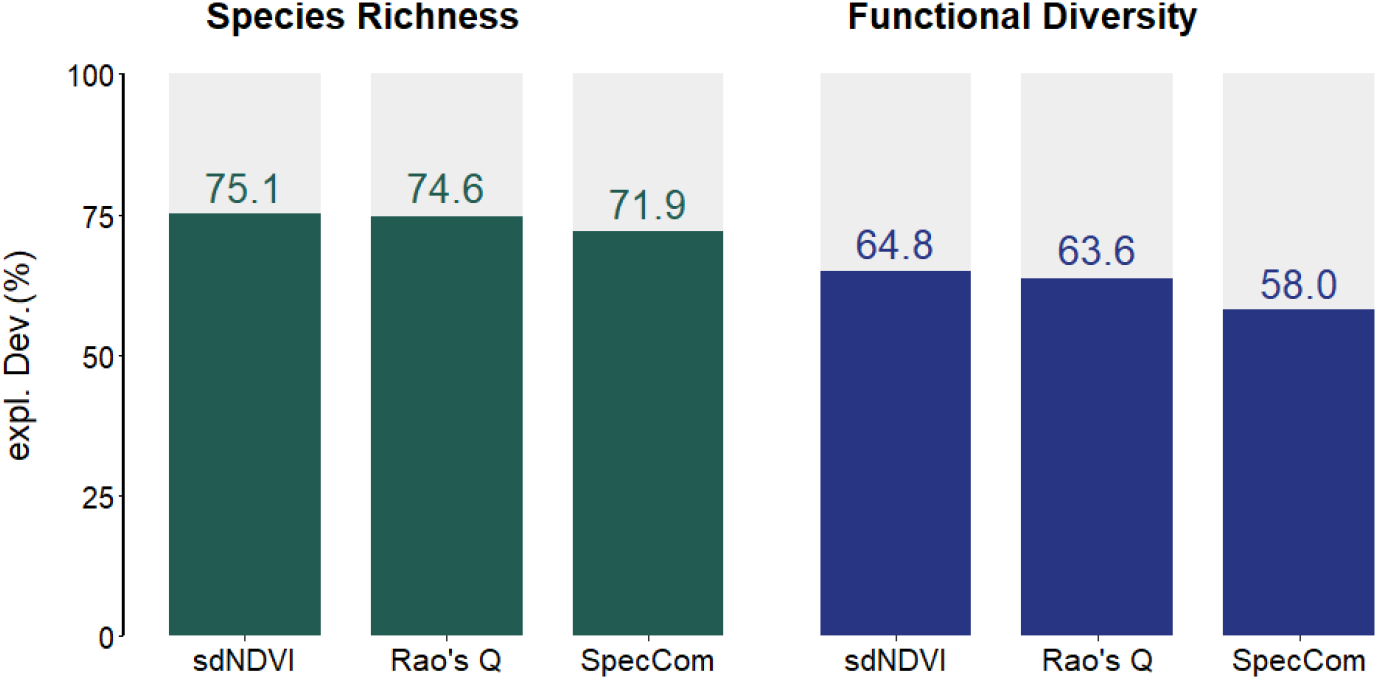
Total explained percentage deviance of models explaining species richness and functional diversity with three types of spectral diversity (sdNDVI, Rao’s Q, SpecCom), sampling effort, and spatial smoothing splines

We found slight differences in the deviance explained by the models among the performances of the SD metrics considered (Figure 2). For both species richness and functional diversity, the models where SD was calculated using SpecCom showed the lowest explained deviance, while the models with continuous metrics (i.e., sdNDVI and Rao’s Q) showed the highest explained deviance.

### 3.1 Variation partitioning

Variation partitioning showed that the unique contribution of SD accounts for only a small fraction of the total variability, ranging from 2.6% to 5.8% in SR models (Figure 3a, c, and e), and from 5.7% to 12.5% in FD models (Figure 3b, d, and f). Moreover, in SR models, sampling effort alone explained almost half of the deviance (between 46.1% and 47.7%) (Figure 3a, c, and e). In contrast, in FD models, most of the variability was jointly explained by SD and splines (between 25.5% and 29.6%) and by spatial autocorrelation (splines) alone (between 22.5% and 26.6%) (Figure 3b, d, and f).

**Figure 3.**
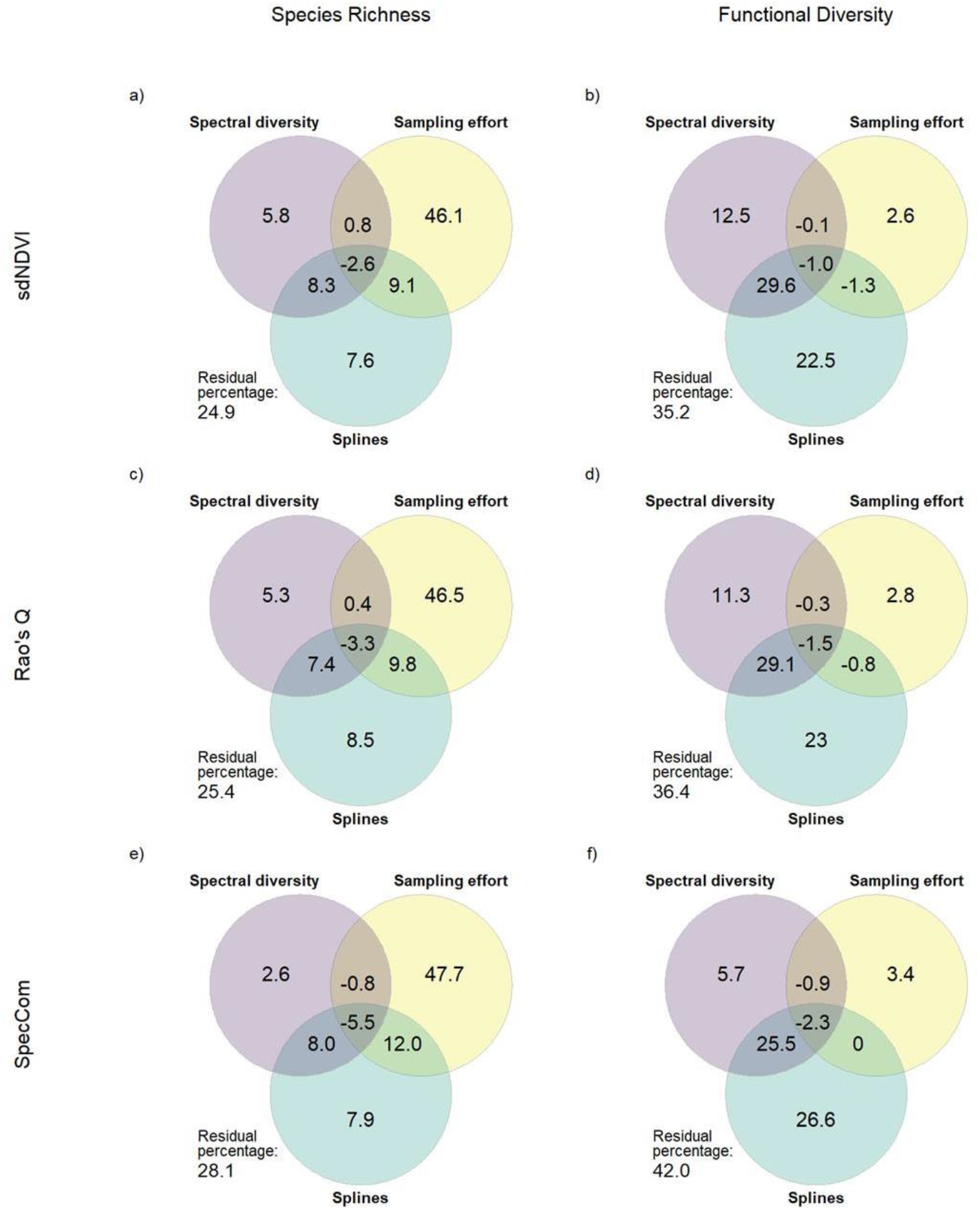
Venn diagrams showing the percentage of explained deviance in plant diversity explained by the three groups of predictors: spectral diversity, sampling effort, and spatial autocorrelation (splines). Overlaps among circles represent fractions of the variation that cannot be unequivocally attributed to one of the overlapping predictors. For example, when spectral diversity is also spatially autocorrelated, some fraction of the explained deviance will be shared between splines and spectral diversity. Negative numbers can arise due to the spline optimisation algorithm during model fitting, are usually small, and can be interpreted as zeroes.

### 3.2 Effect of habitat type

For simplicity, we focus primarily on how the direction of the relationship between SD and biodiversity varies with the different CLC types, therefore we describe only the effects of the interaction between the first power of the SD variables and ecosystem type, since it is the power that determines the direction of the effect (the full list of coefficient estimates can be found in Appendix C). We found that the significance, magnitude, and sign of the effect of SD varied depending on the CLC type dominating the landscape (i.e., grid cell) (Figure 3). Specifically, in the landscapes dominated by arable lands (CLC 21), coniferous forests (CLC 312), and transitional woodland-shrub (CLC 324), the effect of SD was always significant and positive (Figure 4a, b, c, d, e, and f). In addition, the effects of SD in transitional woodland-shrubs (CLC 324) were remarkably high compared to the other interactions (Figure 4), suggesting that increases in SD were associated with a higher increase in plant diversity in transitional woodland-shrubs than in other habitat types. In the case of artificial surfaces (CLC 1), the effect of SD was significant and positive only when considering sdNDVI and Rao’s Q in both SR and FD models (Figure 4a, b, c, and d). The effect in pastures (CLC 231) was significant and positive in all FD models and in the SR model where SD was calculated through sdNDVI (Figure 4a, b, d, and f). The effect in broad-leaved forests (CLC 311) was always significantly positive, except in the SR model where SD was calculated using Rao’s Q (Figure 4c). The effect in mixed forests was, instead, significant and positive only in the SR model where SD was calculated through SpecCom (Figure 4e). The only case where we observed a significant negative effect of the interaction with the predominant habitat type was when considering artificial surfaces (CLC 1) in the models relating plant diversity to SpecCom (Figure 4e and f). In all other cases, the predominant habitat type did not affect the relationship between SD and plant diversity, as the results were not significant.

**Figure 4.**
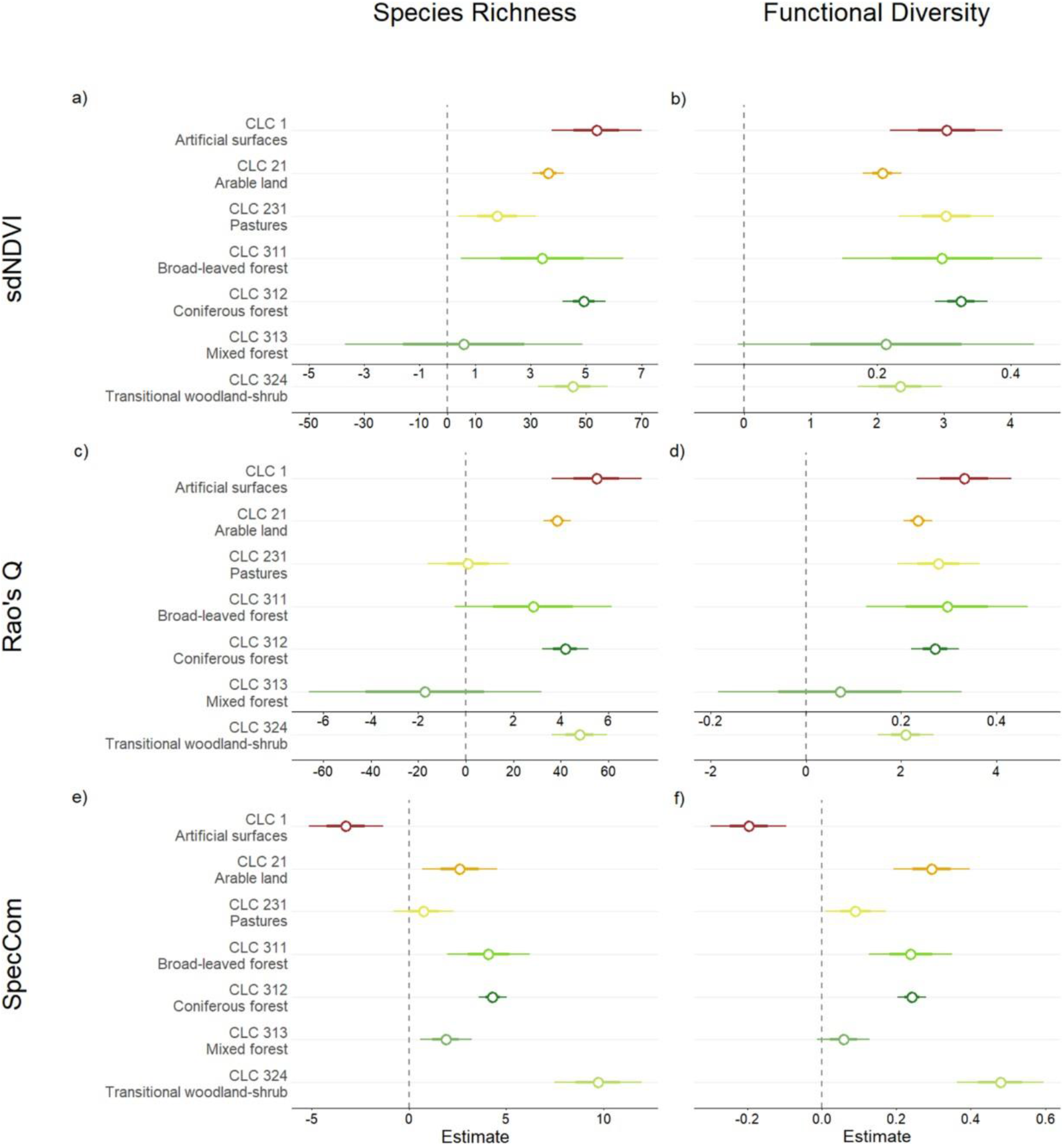
Effect sizes (standardised coefficients) of interactions between SD and land-cover types in the plant diversity ∼ SD models. The dots represent coefficient estimates; the outer error bars are 95% confidence intervals, and the inner error bars indicate one standard error on each side. When the 95% coefficient interval includes a value of zero, the interaction between SD and land-cover type is not significant. Due to the different value ranges of CLC 324 estimates, a second x-axis with appropriate scaling is shown.

## 4. Discussion

### 4.1 Spectral diversity and plant diversity

When modelling the ability of different types of SD metrics to explain the variation of plant diversity at the landscape scale, despite the relatively high goodness-of-fit (percentage of explained deviance) observed in our models, only a small fraction of the variation was explained by SD alone. SD accounted for a larger fraction of variation in FD models, although SR models had a better goodness-of-fit. These results suggest that, even at this scale, SD is linked to functional diversity slightly better than to species richness. Nevertheless, SD and splines combined share a relatively large fraction of variation in all models, suggesting that it is difficult to disentangle the influence of spatial autocorrelation of SD from the effect of SD itself. Overall, the unique contributions of sampling effort and splines explain a large portion of the variation in SR and FD models, respectively. Therefore, routinely inferring plant diversity from SD may lead to inaccurate results, if sampling effort and spatial autocorrelation are not accounted for in situations similar to ours.

Results were generally consistent across species richness and functional diversity, suggesting that SD is able to reflect both aspects of plant diversity at the given landscape scale, which has been reported in previous studies (Frye et al., 2021). This is in agreement with the original formulation of the SVH (Palmer et al., 2002): the heterogeneity of the landscape estimated through SD reflects the diversity of available niches and thus both taxonomic and functional diversity. Indeed, SD does not attempt to directly estimate any specific aspect of plant biodiversity (e.g., taxonomic, phylogenetic, functional), but encompasses them all through physical and ecological rules (Laliberté et al., 2020; Schweiger et al., 2018; Wang and Gamon, 2019).

### 4.2 Spectral diversity and effects of habitat type

The differences found in the relationship between SD and ground diversity across the habitat types considered were based mainly on the ability of the different metrics used (i.e., continuous and categorical) to capture the relationship and on the strength of the relationship.

In landscapes dominated by artificial surfaces, the ability of continuous metrics of SD to predict biodiversity unexpectedly improved. This is because, unlike categorical spectral metrics, continuous metrics are sensitive to extreme reflectance values of background material (e.g., bare soil, litter, rocks), which can have a considerable influence on the SD estimation (Fassnacht et al., 2022; Wang et al., 2018b). At finer resolutions, this issue can be tackled by masking out soil pixels and unvegetated areas (e.g., Gholizadeh et al., 2018). At coarser spatial resolutions, as in our case, such masking may result in the erroneous filtering of mixed pixels corresponding to scarcely vegetated - but biologically relevant – areas, resulting in a consequent loss of information (Schmidtlein and Fassnacht, 2017). Landscape mosaics dominated by urban areas are likely to be very species-rich, as semi-natural vegetation within or adjacent to cities often harbours more plant species than the surrounding landscape (Araújo, 2003; Kühn et al., 2004). Moreover, masking landscape features based on their reflectance values would contradict the original formulation of SVH, which states that it is the magnitude of variation in spectral characteristics of an area that relates to habitat (or vegetation type) heterogeneity and thus to available niche space. In contrast, in landscapes dominated by urban habitats, the ability of the categorical approach to make inferences about biodiversity was impaired, which suggests that these metrics do not properly capture the heterogeneity of this landscape type

The categorical approach of the SpecCom metric can still offer relevant benefits (Fassnacht et al., 2022; Schmidtlein and Fassnacht, 2017). Estimating SD by means of spectral type classification allows summarising the continuity of spectral space into discrete spatial objects that are likely to correspond to distinct landscape features. With this approach, there is less risk of disproportionate influence of extreme pixel values on the spectral variation metric, since they would constitute individual categories among equally meaningful others (Wang and Gamon, 2019). Moreover, spectrally homogeneous spatial units can be identified even when plant optical characteristics change throughout the year according to species phenology, allowing for temporal consistency in spectral type classification. However, this is unlikely to work in larger regions with diverse environmental conditions and, thus, non-synchronous phenological states across similar ecological entities (e.g., communities). In landscapes dominated by transitional woodland-shrub habitats, both categorical and continuous SD metrics exhibited the strongest (albeit still weak) association with plant diversity. Given the inherent spatial vegetation heterogeneity of such areas, which can likely be detected through RS imagery, these results are unsurprising. Indeed, the habitat type is characterised by patchy, bushy and herbaceous vegetation with occasional scattered trees (Bossard et al. 2000), and the resulting spectral heterogeneity can be detected even by current, non-commercial spaceborne sensors.

In landscapes dominated by arable land and coniferous forests, we found the same general positive (although weaker) effect on the relationship between SD and diversity, indifferently of which metric was used. The agreement among SD metrics indicates that the relationship between SD and ground biodiversity is significant and stable. The landscapes dominated by arable land are widespread in Central Europe (as in the case of the Czech Republic, see Appendix B) and are characterised by high spectral heterogeneity, mostly given by the mosaic of heterogeneous and species-poor cultivation patterns. However, these landscapes may harbour several small “islands” of natural and semi-natural species-rich habitats (e.g., grasslands and shrublands), (Wang et al., 2018a) which are captured by SD. In the case of landscapes dominated by coniferous forests, the low spectral heterogeneity of the canopy is coupled with low plant diversity, which is mirrored by the positive SD-diversity relationship.

In our study area, areas classified as pastures (CLC 231) correspond to managed grasslands and meadows. Therefore, the positive and significant interaction between pastures as the predominant habitat type and SD, observed in FD models and SR model using sdNDVI, was less expected than for other habitat types. Indeed, the wide mismatch between pixel resolution and size of individuals and populations of plant species has no negative effect on the SD-diversity relationship, as would be expected based on the smoothed reflectance signal (Wang et al., 2018a; Fassnacht et al., 2022).

### 4.3 Choice of spectral diversity metrics

We found that all our metrics of SD can explain a comparable proportion of the variability in the ground diversity at the scale applied. However, there were differences between categorical (i.e., SpecCom) vs continuous metrics (i.e., Rao’s Q and sdNDVI), with the continuous metrics performing slightly better. Their performance differed depending on the habitat type considered.

The SD metrics that we used all performed similarly, despite differences within different habitat types. However, some practical aspects could be crucial in the choice of the most appropriate metric.

The application of categorical metrics for the routine production of RS biodiversity products with global consistency is complicated by the need for the user’s input. Indeed, it is necessary to conduct preliminary experiments to find the perfect compromise between the number of clusters and the computational effort that would allow reliable results (Féret and de Boissieu, 2020; Rocchini et al., 2021). Regarding continuous metrics, those used in this study differ in the spectral information used to compute SD. Indeed, metrics based on variation in vegetation indices (e.g., sdNDVI) rely on the information provided by specific wavebands that are related to vegetation condition and biomass, and allow discrimination between vegetated and non-vegetated areas, and between different vegetation types. However, the use of this specific (limited) set of spectral bands may exclude some important information about other environmental and biochemical properties (e.g., leaf water content, nitrogen content, pigments, and lignin) (Asner and Martin, 2009). In contrast, spectral entropy metrics (e.g., Rao’s Q) use all the available spectral information and describe the spectral “dimensionality” of the data (Wang and Gamon, 2019), but they can be computationally costly, possibly needing several days to be computed for large areas at a medium-high spatial resolution. As we showed that a simpler vegetation index-based metric (sdNDVI) performs comparably well to a spectral entropy-based Rao’s Q, we recommend using the simpler and computationally low-cost sdNDVI, which already proved performing in other studies (Gholizadeh et al., 2019). However, since previous results (Gholizadeh et al., 2018; Wang et al., 2018b) suggest that the most informative spectral regions for biodiversity assessment can vary with spatial resolution, data properties, and scale of observation, further analyses would be needed in order to assess whether this metric should be used in other analytical contexts.

## 5. Conclusions

We examined the relationship between vascular plant diversity and spectral diversity across more than 2000 grid cells covering the Czech Republic, taking into account the effects of land cover. We investigated the potential of three different SD metrics to infer both species richness and functional diversity. Our results show little to no difference among the metrics tested at this scale, indicating that the association between SD and plant diversity is significant and stable. However, we cannot confirm the general validity of the SVH. Indeed, we showed that in our setting, only a small percentage of variation could be explained solely by SD and that not accounting for sampling effort and spatial autocorrelation in the models could be misleading. In addition, we found an obvious effect of the habitat type prevalent in the landscape on the relationship between SD and biodiversity, which is strongest in areas with transitional habitats between forest and shrubs.

Despite the observed significance of the relationship between SD and ground plant diversity, the strong context-dependence of such a relationship suggests that users should always account for the contextual applicability of SD for mapping biodiversity across space. Along with spatial variation in biodiversity, there is room for testing if the information conveyed by SD could be efficiently used to monitor biodiversity changes over time, e.g., by assessing landscape changes based on the remotely-sensed spectral signal.

Although the SD-biodiversity relationship was not strong compared to the other predictors analyzed, it was still highly significant. Thus, SD has a low potential to serve as the one single proxy for species or functional diversity. However, it still has potential to improve models and predictions of taxonomic or functional diversity. We suggest that, for this purpose, SD should be used as one of several predictors, alongside other well-known variables that affect diversity, such as productivity, climate, and historical processes (Ricklefs & Schluter 1994; Lomolino et al. 2017; Hawkins et al. 2003).

## Acknowledgements

We thank Vojtěch Barták for useful insights and suggestions that greatly served this study.

MP was supported by IGA (Internal Grant Agency) grant from the Faculty of Environmental Sciences at Czech University of Life Sciences (project number 2021B0010). JM and JW were supported by RVO 67985939 (Czech Academy of Sciences). PK was supported by REES (Research Excellence in Environmental Science) grant from the Faculty of Environmental Sciences at Czech University of Life Sciences. JD was supported by the Technology Agency of the Czech Republic (project no. SS02030018) and by RVO 68145535 (Czech Academy of Sciences). MC was supported by the Czech Science Foundation (project no. 19-28491X).

## Appendix A

**Table A1.**
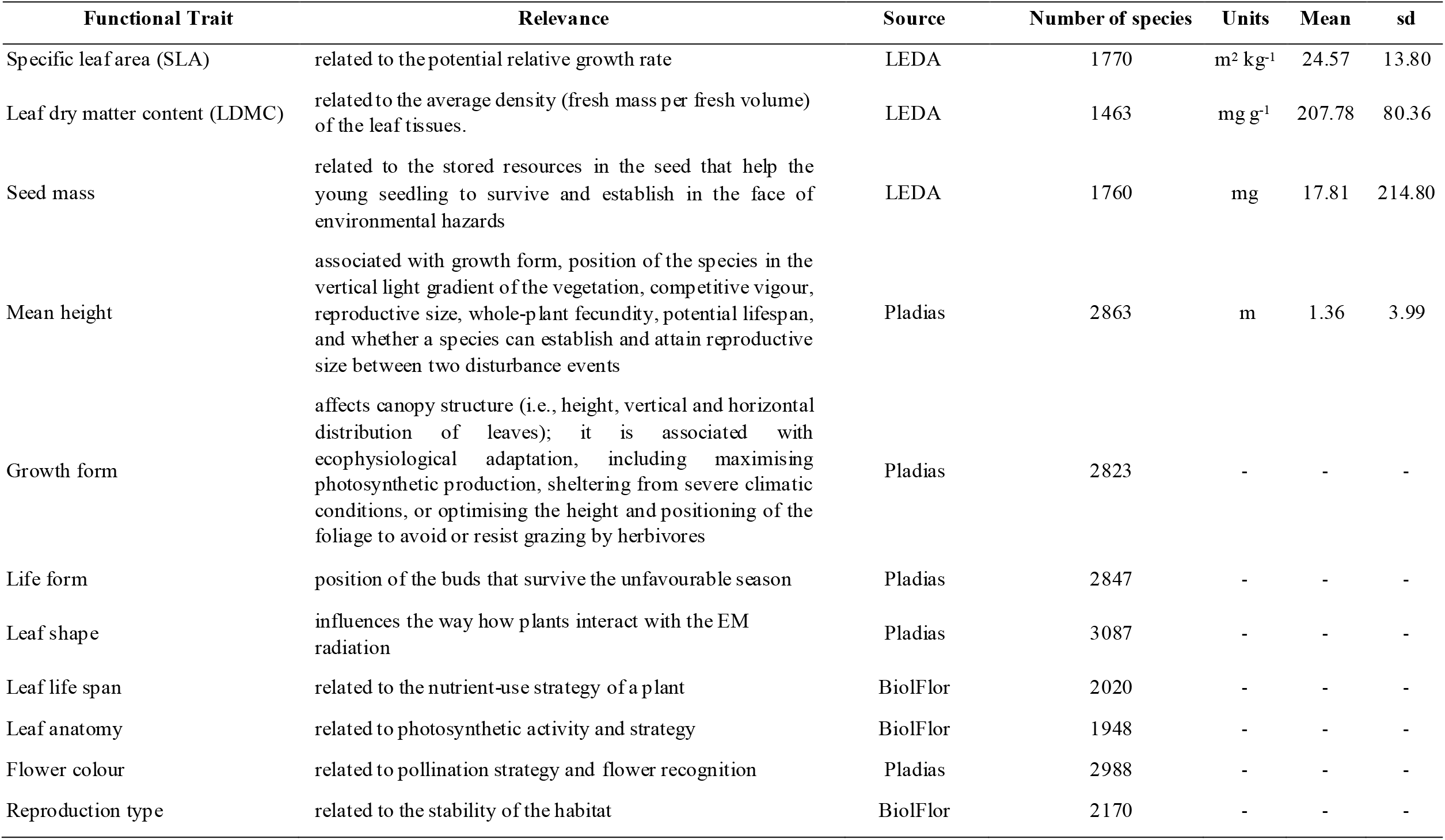
Functional traits used to calculate functional diversity (with their source database) (Chytrý et al., 2021; Pérez-Harguindeguy et al., 2013; Raunkiaer, 1934).

| Functional Trait | Relevance | Source | Number of species | Units | Mean | sd |
| --- | --- | --- | --- | --- | --- | --- |
| Specific leaf area (SLA) | related to the potential relative growth rate | LEDA | 1770 | m<sup>2</sup> kg<sup>-1</sup> | 24.57 | 13.80 |
| Leaf dry matter content (LDMC) | related to the average density (fresh mass per fresh volume) of the leaf tissues. | LEDA | 1463 | mg g<sup>-1</sup> | 207.78 | 80.36 |
| Seed mass | related to the stored resources in the seed that help the young seedling to survive and establish in the face of environmental hazards | LEDA | 1760 | mg | 17.81 | 214.80 |
| Mean height | associated with growth form, position of the species in the vertical light gradient of the vegetation, competitive vigour, reproductive size, whole-plant fecundity, potential lifespan, and whether a species can establish and attain reproductive size between two disturbance events | Pladias | 2863 | m | 1.36 | 3.99 |
| Growth form | affects canopy structure (i.e., height, vertical and horizontal distribution of leaves); it is associated with ecophysiological adaptation, including maximising photosynthetic production, sheltering from severe climatic conditions, or optimising the height and positioning of the foliage to avoid or resist grazing by herbivores | Pladias | 2823 | - | - | - |
| Life form | position of the buds that survive the unfavourable season | Pladias | 2847 | - | - | - |
| Leaf shape | influences the way how plants interact with the EM radiation | Pladias | 3087 | - | - | - |
| Leaf life span | related to the nutrient-use strategy of a plant | BiolFlor | 2020 | - | - | - |
| Leaf anatomy | related to photosynthetic activity and strategy | BiolFlor | 1948 | - | - | - |
| Flower colour | related to pollination strategy and flower recognition | Pladias | 2988 | - | - | - |
| Reproduction type | related to the stability of the habitat | BiolFlor | 2170 | - | - | - |

## Appendix B

**Figure B1.**
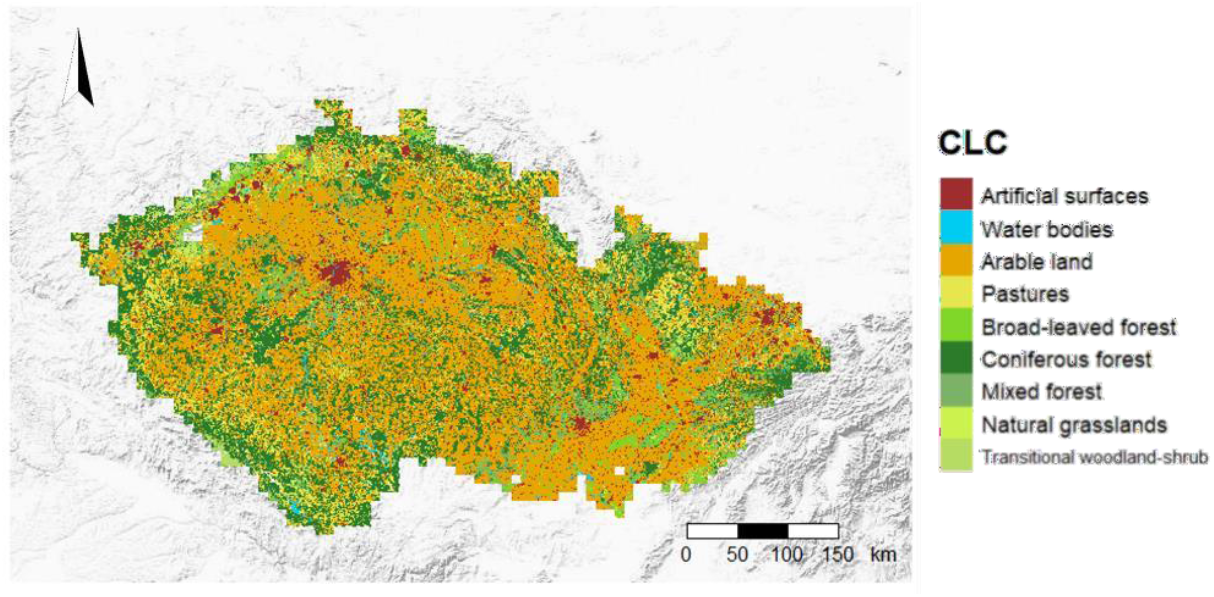
Corine Land Cover map of the study area (reference year 2012, 100 m resolution).

**Figure B2.**
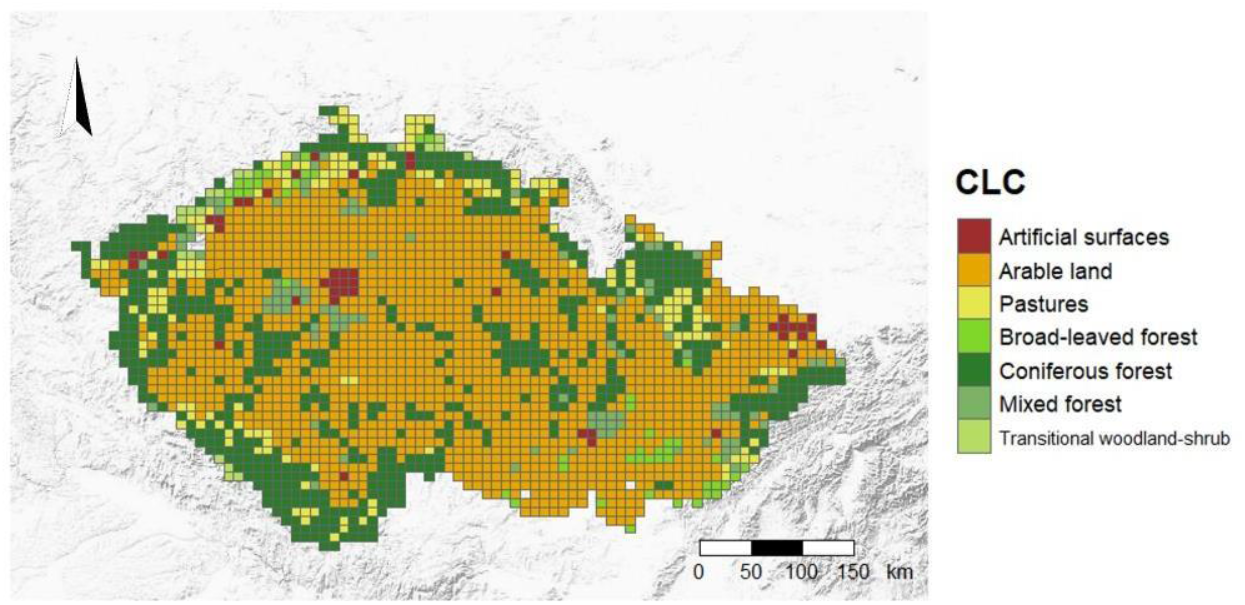
Map of the most abundant CLC type within each grid cell.

**Table B1.** Legend of the land cover codes used, with their respective level of detail within the CLC classification and number of grid cells.

| Code | CLC Level | Cover type | N |
| --- | --- | --- | --- |
| 1 | I | Artificial surfaces | 46 |
| 21 | II | Arable land | 1381 |
| 231 | III | Pastures | 161 |
| 311 | III | Broad-leaved forest | 49 |
| 312 | III | Coniferous forest | 608 |
| 313 | III | Mixed forest | 100 |
| 321 | III | Natural grasslands | 3 |
| 324 | III | Transitional woodland-shrub | 18 |

## Appendix C

**Table C1.** Coefficient estimate values of the interaction between SD and land-cover type in the plant diversity ∼ SD models. Significance codes: *** (p-value < 0.001), ** (p-value < 0.01), * (p-value < 0.05).

|  | Species Richness |  |  |  |  |  | Functional Diversity |  |  |  |  |  |
| --- | --- | --- | --- | --- | --- | --- | --- | --- | --- | --- | --- | --- |
|  | sdNDVI |  | Rao's Q |  | SpecCom |  | sdNDVI |  | Rao's Q |  | SpecCom |  |
| CLC 1 1<sup>st</sup> power | 5.38 | *** | 5.50 | *** | -3.23 | *** | 0.30 | *** | 0.33 | *** | -0.20 | *** |
| CLC 1 2<sup>nd</sup> power | -1.85 | * | -1.58 | * | -1.59 |  | -0.07 |  | -0.09 | * | 0.01 |  |
| CLC 21 1<sup>st</sup> power | 3.64 | *** | 3.85 | *** | 2.61 | ** | 0.21 | *** | 0.23 | *** | 0.30 | *** |
| CLC 21 2<sup>nd</sup> power | -1.18 | *** | -1.18 | *** | 3.16 | * | -0.10 | *** | -0.10 | *** | 0.15 | * |
| CLC 231 1<sup>st</sup> power | 1.79 | * | 0.09 |  | 0.76 |  | 0.30 | *** | 0.28 | *** | 0.09 | * |
| CLC 231 2<sup>nd</sup> power | -3.26 | *** | -3.69 | *** | 1.88 |  | -0.06 | * | -0.03 |  | 0.08 |  |
| CLC 311 1<sup>st</sup> power | 3.42 | * | 2.83 |  | 4.09 | *** | 0.30 | *** | 0.30 | *** | 0.24 | *** |
| CLC 311 2<sup>nd</sup> power | -3.64 |  | -3.15 |  | 2.09 |  | -0.17 |  | -0.13 |  | 0.09 |  |
| CLC 312 1<sup>st</sup> power | 4.92 | *** | 4.18 | *** | 4.29 | *** | 0.33 | *** | 0.27 | *** | 0.24 | *** |
| CLC 312 2<sup>nd</sup> power | -2.68 | *** | -2.65 | *** | 0.79 | *** | -0.18 | *** | -0.19 | *** | 0.06 | ** |
| CLC 313 1<sup>st</sup> power | 0.58 |  | -1.71 |  | 1.90 | ** | 0.21 |  | 0.07 |  | 0.06 |  |
| CLC 313 2<sup>nd</sup> power | -4.66 | * | -5.38 | * | 0.03 |  | -0.01 |  | -0.11 |  | 0.00 |  |
| CLC 324 1<sup>st</sup> power | 45.28 | *** | 47.80 | *** | 9.70 | *** | 2.34 | *** | 2.09 | *** | 0.48 | *** |
| CLC 324 2<sup>nd</sup> power | 26.51 | *** | 31.34 | *** | 3.86 | *** | 1.43 | *** | 1.26 | *** | 0.17 | *** |

